# Cyclic di-AMP inhibits *Listeria monocytogenes* thymineless death during infection

**DOI:** 10.1101/2025.05.14.654034

**Authors:** Joshua P. Leeming, Omar M Elkassih, Damilola T. Oyebode, Joshua J. Woodward, Qing Tang

**Author notes:** Address correspondence to Qing Tang. Joshua P. Leeming and Omar M Elkassih contributed equally to this work. Author order was determined on the basis of seniority.

## Abstract

Anti-folate antibiotics are used to treat meningitis and refractory listeriosis caused by drug-resistant *Listeria monocytogenes* (*Lm*). Their bactericidal activity is attributed to the deactivation of thymidylate synthase (ThyA), which subsequently induces bacterial cell death when thymidine is depleted—a process known as thymineless death (TLD). Despite decades of study, the mechanisms of TLD, especially during infection, remain unclear. Cyclic di-AMP (c-di-AMP), a common bacterial second messenger that regulates bacterial stress responses, is elevated in response to anti-folate antibiotics. In this study, we found that elevated c-di-AMP is required to inhibit TLD in *Lm*. Conversely, reducing c-di-AMP levels in the Δ*thyA* mutant led to increased bacterial cell death under thymidine starvation and significant reduction in intracellular growth. Furthermore, we found that Δ*thyA* exhibited a more pronounced growth defect during oral infection compared to intravenous infection, due to limited thymidine availability in the gallbladder, which acts as a bottleneck for Δ*thyA* in establishing infection. Notably, decreasing c-di-AMP levels abolished the infection capacity of Δ*thyA* in both infection models. Finally, we identified that the c-di-AMP-binding protein PstA contributes to bacterial cell death when c-di-AMP concentrations are low. Deletion of *pstA* in the Δ*thyA* background rescued the elevated cell death caused by c-di-AMP depletion both *in vitro* and during mouse infections. Our study identifies a previously unrecognized mechanism of TLD regulation mediated by c-di-AMP. This expands fundamental knowledge of TLD in the context of infection and provides insight into potential combined therapeutic strategies for listeriosis targeting both anti-folate and c-di-AMP metabolic pathways.

## Introduction

*Listeria monocytogenes* (*Lm*) is a Gram-positive, facultative intracellular pathogen found in natural ecosystems and widely distributed across various environmental niches (1, 2). Consuming food contaminated with *Lm* can lead to severe invasive infections, resulting in a life-threatening disease known as listeriosis (3). Listeriosis commonly presents as bacteremia, meningitis, or meningoencephalitis and includes pregnancy-associated infections, which may lead to miscarriage or neonatal sepsis (4). Listeriosis is a leading cause of death from foodborne illness, with a 20% fatality rate. Nearly all cases require hospitalization, and 25% of pregnancy-related cases result in fetal or neonatal death (5). Antibiotics remain the primary treatment for listeriosis. However, over 30% of patients experience antibiotic therapy failure (6). The underlying reasons for this failure remain unclear, particularly due to the limited understanding of how antibiotics act on the intracellular niches of *Lm* and how the bacterium adapts its physiology in response. This knowledge gap significantly hinders progress in developing optimized therapies for listeriosis.

Anti-folate antibiotics are widely used in clinical practice to treat various bacterial infections. The antifolate antibiotic combination trimethoprim-sulfamethoxazole, also known as co-trimoxazole, has bactericidal activity against both extracellular and intracellular *Lm* and exhibits excellent penetration into various organs, including the central nervous system (CNS). As a result, they are used to treat meningitis and refractory listeriosis caused by drug-resistant *Lm* (7–10). Moreover, antifolate antibiotics are used for individuals who are allergic to penicillins (6, 11). Anti-folate antibiotics function by inducing thymineless death (TLD) of bacteria, a phenomenon in which cells rapidly lose viability due to thymidine starvation. Despite their long-standing clinical use against listeriosis, the mechanism by which antifolates inhibit the growth of *Lm* and how bacteria respond to LTD remains largely unexplored.

TLD was first discovered in *Escherichia coli* over 70 years ago (12), and since then, *E. coli* has become the most widely used model for studying this phenomenon in prokaryotes. Although the exact mechanism of TLD remains unclear, increasing evidence suggests that it involves multiple pathways, including uncoordinated chromosomal replication initiation, central metabolism or protein synthesis (13, 14), DNA damage-response (15), reactive oxygen species (ROS) production (16), intracellular acidification (17), and abnormal cytoplasm and cell envelope (18). Despite these advances, there remain open questions about the mechanisms underlying TLD in the context of infection. Notably, it is still unclear whether, and how, bacteria acquire thymidine from the host environment. *Lm* has emerged as a promising model for studying TLD during infection due to its sensitivity to antifolate antibiotics and its capacity for intracellular replication.

In response to external antibiotic stresses, *Lm* utilizes various mechanisms to rapidly and precisely alter its bacterial physiology, facilitating its survival. One such mechanism is the global regulation of bacterial physiology by the second messenger cyclic di-AMP (c-di-AMP). C-di-AMP is synthesized from two ATP molecules by diadenylate cyclase (DacA), and it can be degraded to pApA, or two AMP molecules, by specific phosphodiesterases GdpP or PdeA, respectively (19, 20). Additionally, intracellular c-di-AMP concentrations of *Lm* are regulated by multidrug transporters (MdrM) that efflux this nucleotide (20). Through binding to RNA or protein regulators, c-di-AMP regulates cell wall integrity (21), osmotic homeostasis (22), and central metabolism (23). In *Lm*, both β-lactam and antifolate antibiotics have been shown to enhance c-di-AMP production (24), indicating the importance of c-di-AMP in regulating bacterial fitness against these antibiotics. This is further supported by multiple studies demonstrating that c-di-AMP level positively correlates with resistance to β-lactam antibiotics in several human pathogens (25–33). It remains unclear whether c-di-AMP functions as a regulator of antifolate antibiotic susceptibility and the underlying mechanisms involved.

In this study, we identified that c-di-AMP is a global regulator of antibiotic susceptibility to β-lactams, antifolates, and multiple antibiotics targeting protein synthesis. Our previous work showed that deactivation of ThyA, either through antifolate treatment or deletion of the thyA gene (Δ*thyA*), leads to elevated levels of c-di-AMP production (24). In this study, we further demonstrate that the elevated c-di-AMP is necessary to inhibit TLD of *Lm* Δ*thyA*. Furthermore, we found that eukaryotic host cells do not provide a sufficient thymidine source to support the intracellular growth of Δ*thyA*, which ultimately leads to a growth defect during systemic infection. We also identified that the conserved c-di-AMP-binding protein, PstA, contributes to cell death of Δ*thyA* under low c-di-AMP conditions. By investigating the role of c-di-AMP in regulating TLD during infection, our study reveals a previously unrecognized mechanism of bacterial TLD and expands fundamental knowledge of c-di-AMP signaling in cellular stress responses, offering potential therapeutic insights targeting this pathway.

## Results

### C-di-AMP regulates antifolate susceptibility of *Lm*

To investigate whether c-di-AMP regulates antibiotic susceptibility of *Lm*, the Δ*dacA*::*disA* strain—where DacA is replaced by an IPTG-inducible DisA (*Bacillus subtilis* diadenylate cyclase (34))—was cultured without IPTG to halt c-di-AMP production. C-di-AMP regulates various processes including osmolyte transport and bacterial cell wall integrity. These regulations subsequently affects bacterial susceptibility to β-lactam antibiotics in *S. aureus* (25, 26, 28, 29), *Lm* (21, 32), *Streptococcus pyogenes* (27, 31) and *Enterococcus faecalis* (35). Consistent with these studies, the Δ*dacA*::*disA* strain displayed increased susceptibility to β-lactam antibiotics compared to the wild-type (WT) strain. Furthermore, the low c-di-AMP strain is susceptible to various categories of protein synthesis-inhibiting antibiotics, indicating the involvement of complex mechanisms. It also exhibited heightened susceptibility to most of the tested antifolate antibiotics (**Fig. 1A**). This phenotype was further validated through disk diffusion assays, where treatment with antifolates, including SXT (<u>S</u>ulfametho<u>x</u>azole-<u>T</u>rimethoprim), Sulfamethoxazole, and Trimethoprim, resulted in a larger inhibition zone on plates containing the Δ*dacA*::*disA* strain without IPTG induction, compared to both the Δ*dacA*::*disA* strain supplemented with IPTG and the WT strains (**Fig. 1B**). In contrast, all strains exhibited similar susceptibility to the control aminoglycoside antibiotic kanamycin (**Fig. 1C**).

**Figure 1.**
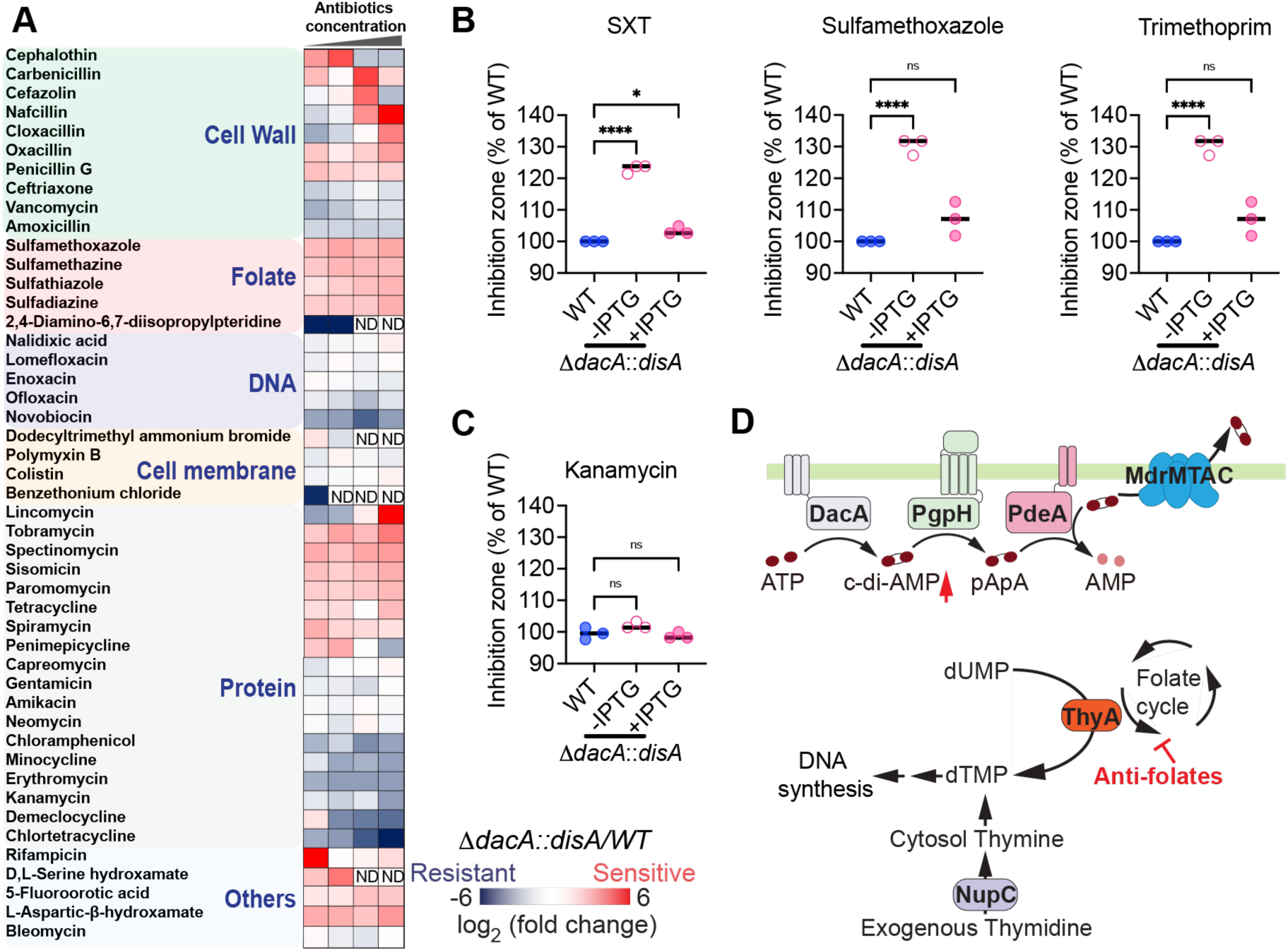
C-di-AMP functions as a global regulator for antibiotic resistance. (A) The heatmap indicates the log_2_ ratio of the CFU recovery of Δ*dacA::disA* (low c-di-AMP strain) relative to WT strain exposed to various antibiotics. The Δ*dacA::disA* strain, in which DacA was replaced by an IPTG-inducible diadenylate cyclase, was cultured without IPTG for 16 h to halt c-di-AMP production. Blue texts indicate antibiotic classes according to their mechanisms of action, “ND” indicates that no bacteria recovery after treatment and antibiotic susceptibility was not determined. (B) and (C) Antibiotic susceptibility of *Lm* strains measured by disk diffusion assays. The inhibition zone of each strain was normalized to that of the WT. For all panels, mean values of biological replicates are plotted, and error bars indicate ±SD*, P* values were calculated using Ordinary one-way analysis. Asterisks indicate that differences are statistically significant (*, *P* < 0.05; ****, *P* < 0.0001), and ns indicates no significant difference. (D) Schematic of thymidine and c-di-AMP metabolism in bacteria. ThyA, thymidylate synthase; NupC, nucleoside permease; DacA, diadenylate cyclase; PgpH, phosphodiesterase; PdeA, phosphodiesterase; MdrMTAC, multidrug transporters.

Anti-folates disrupt the synthesis of 5,10-methylenetetrahydrofolate, an essential co-factor of thymidylate synthase (ThyA). ThyA synthesizes the DNA synthesis precursor dTMP from dUMP. Disruption of the folate cycle inhibits ThyA’s enzymatic activity, resulting in reliance on exogenous thymidine to maintain DNA synthesis (**Fig. 1D**). When deprived of thymidine, ThyA defective strains lose viability, known as TLD. Inactivating ThyA by antifolate inhibition or *thyA* gene deletion leads to elevated c-di-AMP production in *Lm* (24) (**Fig. 1D**), suggesting that *Lm* may increase c-di-AMP production under thymidine deprivation to inhibit TLD.

### C-di-AMP is crucial for maintaining cell viability during thymidine deprivation *in vitro*

Given that inactivation of ThyA by deleting its encoding gene is equivalent to inhibiting ThyA with anti- folates, we generated a *Lm thyA* knockout strain (Δ*thyA*) to study the mechanism of TLD. *Lm* Δ*thyA* produces significantly higher levels of c-di-AMP compared to the WT strain (24). C-di-AMP depletion is detrimental to the growth of *Lm* (32). As a result, we generated WT::*pdeA* and Δ*thyA*::*pdeA* strains, in which we overexpressed the phosphodiesterase PdeA to reduce, but not completely eliminate, c-di-AMP production (**Fig. 2A**). We found that the Δ*thyA*::*pdeA* strain exhibited only a moderate growth delay compared to the Δ*thyA* strain when supplemented with high concentration of exogenous thymidine (10 µg/ml) (**Fig. 2B**, left); however, the growth defect was exacerbated under lower thymidine concentration (1.25 µg/ml) (**Fig. 2B**, right). In contrast, in the WT strain with functional ThyA, decreasing c-di-AMP did not affect bacterial growth regardless of thymidine supplementation (**Fig. 2B**). Additionally, we assessed the cell death of Δ*thyA* and Δ*thyA*::*pdeA* strains under thymidine starvation using live/dead cell staining with propidium iodide. Δ*thyA*::*pdeA* exhibited significantly higher cell death compared to the Δ*thyA* strain after 11 hours of thymidine starvation (**Fig. 2C and Supplementary Video 1-2**). Furthermore, reducing c- di-AMP levels by overexpressing either the full-length (Δ*thyA*::*pdeA*) or the enzymatic domain of its phosphodiesterase in the Δ*thyA* strain (Δ*thyA*::*pdeA*_84-657_) accelerated cell death. Conversely, overexpression of the enzymatically inactive PdeA mutant (Δ*thyA*::*pdeA*_DHH-AAA_) had no effect on cell viability, as determined by colony-forming unit (CFU) recovery after thymidine starvation (**Fig. 2D**). These data suggest that elevated c-di-AMP production in Δ*thyA* is crucial for maintaining cell viability when cells are deprived of thymidine.

**Figure 2.**
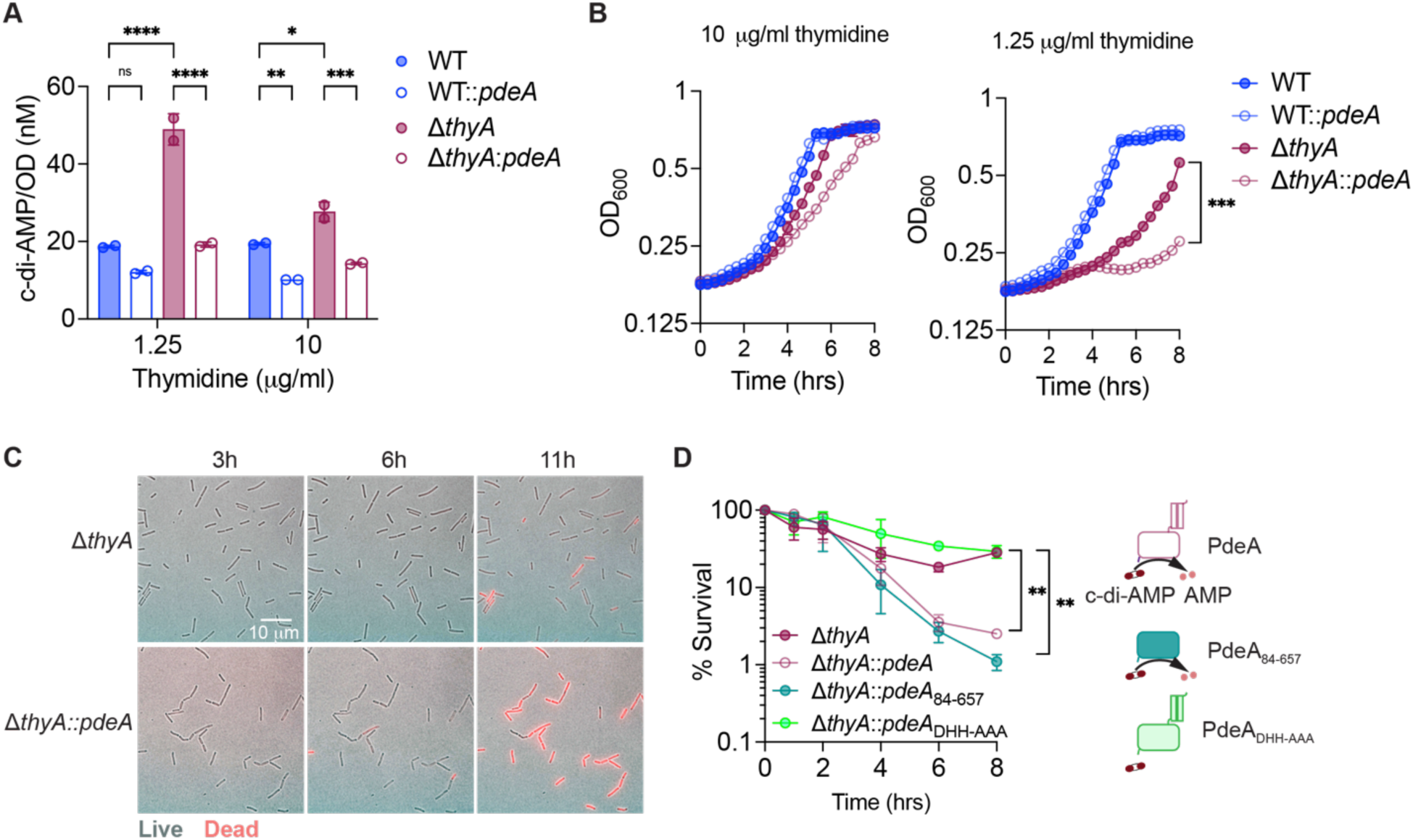
C-di-AMP is crucial for maintaining cell viability during thymidine deprivation *in vitro*. (A) Intracellular c-di-AMP concentration of *Lm* strains quantified using the CDA-Luc assay. The *Lm* strains were grown overnight statically at 30 °C in BHI broth supplemented with different concentrations of thymidine. (B) Growth curves of *Lm* strains with varying concentrations of thymidine supplementation in BHI broth incubated statically at 30 °C. (C) Composite phase-contrast and red fluorescence images of *Lm* strains stained with propidium iodide. Δ*thyA* and Δ*thyA*::*pdeA* strains were grown on LB agar without thymidine and supplemented with propidium iodide during the 11 hours of imaging, which penetrates only the cell envelopes of dead cells. (D) Relative CFU recovery of bacteria incubated in LB broth without thymidine. The percentage of survival at each time point was calculated by normalizing the CFU count to the value at time 0. PdeA, c-di-AMP phosphodiesterase, PdeA_84-657_, enzymatic domain of PdeA. PdeA_DHH- AAA_, enzymatic dead mutant of PdeA. For all panels, mean values of biological duplicates are plotted, and error bars indicate ±SD, *P* values were calculated using either Two-way ANOVA (A) or Ordinary one-way ANOVA (B and D) analysis. Asterisks indicate that differences are statistically significant (*, *P* < 0.05; **, *P* < 0.01; ***, *P* < 0.001, ****, *P* < 0.0001), and ns indicates no significant difference.

### C-di-AMP is required for the intracellular growth of Δ*thyA* in tissue cultures

*Lm* is a highly adaptable bacterium that replicates as a facultative intracellular pathogen. Its intracellular replication within the host is essential for its pathogenesis. However, it remains unclear whether *Lm* can acquire sufficient thymidine from host cells to support its intracellular growth in response to antifolate antibiotics.

To investigate whether c-di-AMP is essential for the intracellular growth of *Lm*, immortalized <u>b</u>one <u>m</u>arrow <u>d</u>erived <u>m</u>acrophages (iBMDMs) were infected with WT, WT::*pdeA*, Δ*thyA*, and Δ*thyA*::*pdeA* strains in the presence or absence of extracellular thymidine (**Fig. 3A**). In the absence of thymidine, Δ*thyA* showed a significant growth defect compared to the WT strain; Notably, Δ*thyA*::*pdeA* totally lost its ability to replicate in iBMDMs (**Fig. 3B**). Nevertheless, supplementation with extracellular thymidine restored the growth defect of both strains (**Fig. 3C**). Additionally, PdeA overexpression in the WT strain did not affect its intracellular growth, with both the WT and WT::*pdeA* strains showing similar growth trends regardless of thymidine supplementation (**Fig. 3B-C**). Moreover, after 6 hours of intracellular growth in macrophages, the WT, WT::*pdeA*, and Δ*thyA* strains exhibited similar bacterial morphology, while Δ*thyA*::*pdeA* cells displayed filamentation (**Fig. 3D-E**). Cell elongation or swelling under thymidine starvation have been observed across multiple bacterial spices (18, 36). Supplementation with exogenous thymidine fully restored the normal morphology of Δ*thyA*::*pdeA* cells (**Fig. 3D-E**). In addition, the elongated Δ*thyA*::*pdeA* cells were able to recruit actin tails, which facilitate intracellular movement and cell-to-cell spread (**Fig. 3D**). These results indicate that Δ*thyA* could not obtain enough thymidine to support its intracellular growth, while c-di-AMP is essential for inhibiting bacterial cell death caused by thymidine deprivation.

**Figure 3.**
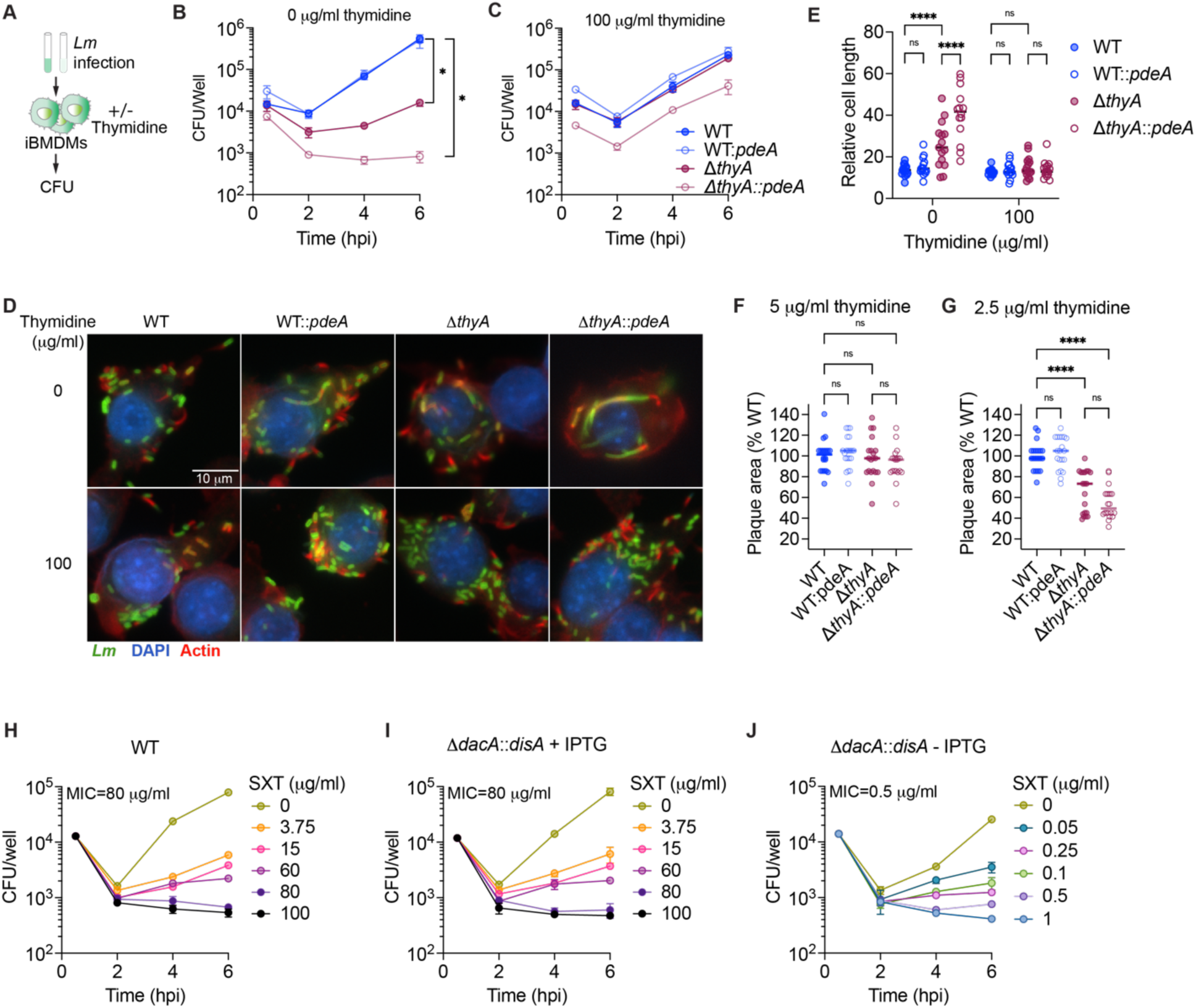
Elevated c-di-AMP production is required for the intracellular growth of Δ*thyA*. (A) Schematic of macrophage infection with *Lm*. The iBMDMs were infected with *Lm* with an MOI of 1 in the presence or absence of extracellular thymidine supplementation. Gentamicin was supplemented at 0.5 hpi to kill the extracellular bacteria. (B) and (C) Intracellular growth curve of *Lm* in iBMDMs as describe in (A). (D) Fluorescence microscopy images of iBMDMs infected with *Lm* at an MOI of 10 at 6 hpi. Nuclei were stained with DAPI (blue), F-actin with Alexa Fluor 568 phalloidin (red), and *Lm* with anti-*Lm* antibody (green). (E) Cell length quantification of intracellular *Lm* as indicated in (D). Mean values of cell length of 15 bacterial cells from three different macrophages were plotted. (F) and (G) Plaque size of L2 fibroblast with *Lm* infection. The monolayer of fibroblasts was infected with *Lm* for 2 days with gentamicin and various concentrations of thymidine supplementation and then stained with neutral red. The plots represent the median of 20 plaques measured from three different wells, with plaque diameters normalized to the control. (H) (I) and (J) Intracellular growth curve of *Lm* in iBMDMs with or without SXT treatment. The iBMDMs were infected with *Lm* with an MOI of 1 in the presence or absence of IPTG supplementation. Gentamicin and SXT was supplemented at 0.5 hpi. For panel (B), (C), (H), and (I), mean values of biological duplicates are plotted. For all panels, error bars indicate ±SD*, P* values were calculated using Ordinary one-way ANOVA (B, C, F, G) or Two-way ANOVA (E) analysis. Asterisks indicate that differences are statistically significant (*, *P* < 0.05; **, *P* < 0.01; ****, *P* < 0.0001), and ns indicates no significant difference.

In addition to macrophages, *Lm* invades and grows in a variety of mammalian cells, including epithelial cells and fibroblasts. *Lm* spreads intracellularly and forms small plaques in monolayers of fibroblasts (37). To determine if c-di-AMP is essential for the growth and intracellular spreading of Δ*thyA*, mouse L2 fibroblasts were infected with *Lm* to measure plaque sizes. Consistent with the defective intracellular growth in macrophages, L2 cells infected with Δ*thyA* did not form any plaques in the absence of exogenous thymine (**Supplementary** Fig. 1). However, supplementation with 5 µg/ml extracellular thymidine fully restored the plaque-forming ability of Δ*thyA* (**Fig. 3F**). Moreover, with a lower concentration of thymidine supplementation (2.5 µg/ml), plaque sizes of both Δ*thyA* and Δ*thyA*::*pdeA* are significantly smaller than those formed by the WT and WT::*pdeA* strains. Δ*thyA*::*pdeA* exhibited a moderate, but not significant, reduction in plaque size compared to Δ*thyA* (**Fig. 3G**). These results indicate that c-di-AMP does not impact the intracellular spreading of Δ*thyA*, as further supported by the observation that Δ*thyA*::*pdeA* forms actin tails (**Fig. 3D**).

Our data indicate that eukaryotic host cells cannot supply sufficient thymidine to support the intracellular growth of Δ*thyA*, and that c-di-AMP is required to inhibits thymineless death. We hypothesize that c-di-AMP is also essential for *Lm* survival during antifolate antibiotic treatment. iBMDMs infected with WT or Δ*dacA*::*disA* strains, with or without IPTG supplementation, were treated with different concentrations of SXT at 0.5 hpi. Consistent with our hypothesis, the minimum inhibitory concentration (MIC) of SXT required to inhibit intracellular growth in iBMDMs is 80 µg/ml for both WT *Lm* and the Δ*dacA*::*disA* strain supplemented with IPTG, whereas the MIC for the Δ*dacA*::*disA* strain without IPTG induction is markedly lower at 0.5 µg/ml (**Fig. 3H-J**). When SXT is used clinically, mean blood levels of SXT range from 98 to 120 μg/ml during the first 8 hours following a single oral dose (38), which is 196 to 240 fold higher than the MIC for Δ*dacA*::*disA* without IPTG. As a result, combination therapy targeting both folate metabolism and c-di-AMP synthesis could significantly enhance the efficacy of antifolate treatments.

### C-di-AMP is required for the growth of Δ*thyA* during mouse infection

To investigate whether c-di-AMP is required by Δ*thyA* to maintain bacterial growth *in vivo*, C57BL/6 mice were retro-orbitally infected with WT, WT::*pdeA*, Δ*thyA*, and Δ*thyA*::*pdeA* strains with an inoculum of 10^5^ CFU/mouse, followed by CFU enumeration of the liver and spleen at 72 hpi (**Fig. 4A**). All strains caused weight loss upon infection; however, the weight of Δ*thyA*::*pdeA*-infected mice stopped decreasing at 48 hpi (**Fig. 4B**), indicating the dampened ability of this strain to cause listeriosis. The Δ*thyA* strain exhibited robust growth in both the liver and spleen, indicating that it had access to thymidine *in vivo* to support its proliferation. Notably, reducing c-di-AMP levels by overexpressing PdeA in the WT strain only slightly impaired its growth, but completely abolished the growth of Δ*thyA* in both the spleen and liver (**Fig. 4C and D**), which is consistent with the reduced weight loss observed in mice infected with Δ*thyA*::*pdeA* (**Fig. 4B**).

**Figure 4.**
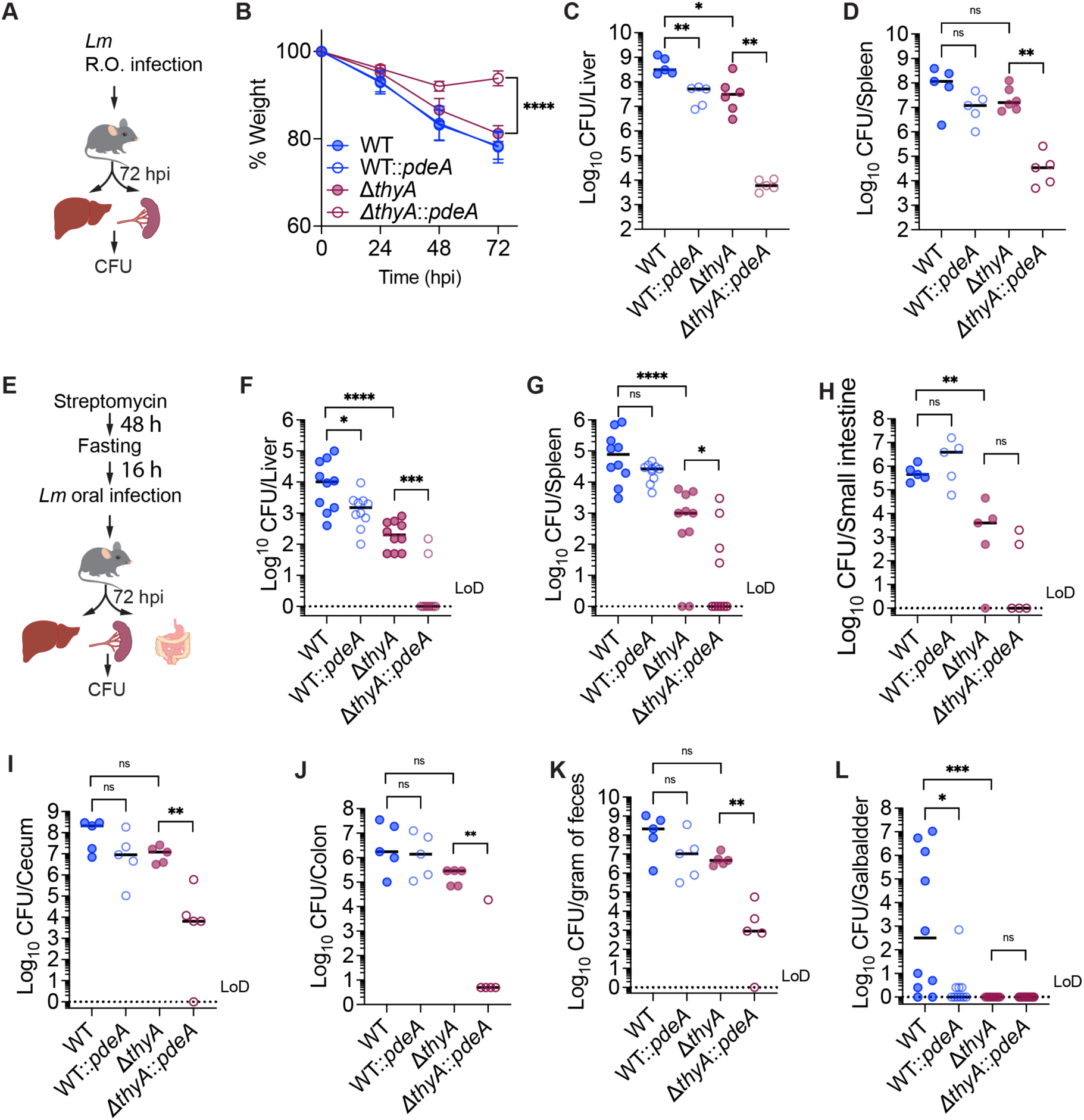
Elevated c-di-AMP production is required for the growth of Δ*thyA* during mouse infection. (A) Schematic of retro-orbital (R.O.) *Lm* infection in mice. C57BL/6 mice were infected with 10^5^ CFU/mouse via the retro-orbital route. Livers and spleens were harvested for CFU enumeration at 72 hpi. (B) Weight loss of infected mice as indicated in panel (A). (C) and (D) CFU recovery of *Lm* from spleens and livers as describe in (A). (E) Schematic of oral *Lm* infection in mice. C57BL/6 mice were given 5 mg/mL streptomycin in drinking water for 48 hours, fasted overnight, and then orally inoculated with 10^8^ *Lm*. Livers and spleens were harvested for CFU enumeration at 72 hpi. (F) to (L) CFU recovery of *Lm* from mouse tissue or feces samples, as described in (E). For panel B, mean values of five biological replicates are plotted, and error bars indicate ±SD*, P* values were calculated using Ordinary one-way ANOVA analysis. For all CFU recovery in mouse infections, biological replicates are plotted. Horizontal black bars indicate the median of the data. *P* values were calculated using Mann-Whitney analysis. Asterisks indicate that differences are statistically significant (*, *P* < 0.05; **, *P* < 0.01; ***, *P* < 0.001, ****, *P* < 0.0001), and ns indicates no significant difference. LoD denotes the limit of detection.

Intravenous *Lm* infection bypasses the intestinal epithelial barrier and directly enters the bloodstream. To determine whether c-di-AMP is required for Δ*thyA* to disseminate from the intestine to peripheral tissues, we orally infected C57BL/6 mice with the same strains with an inoculum of 10^8^ CFU/mouse and monitored the CFU recovery from mouse tissues at 72 hpi (**Fig. 4E**). Δ*thyA* exhibited a more pronounced growth defect during oral infection compared to retro-orbital infection, with approximately a 2-log decrease in CFU recovery from both the liver and spleen compared to the WT. Consistently, we observed that Δ*thyA*::*pdeA* showed an even more severe growth impairment compared to Δ*thyA*, with most mice displaying CFU levels in the liver and spleen below the limit of detection (**Fig. 4F** and **G**). Surprisingly, while Δ*thyA*::*pdeA* showed a pronounced defect in establishing infection in the gastrointestinal tract, Δ*thyA* exhibited comparable or only moderately reduced colonization in the cecum, colon, and feces relative to the WT (**Fig. 4H**-**K**). As gallbladder is an important reservoir for *Lm* during oral infection (39), we found that neither Δ*thyA* nor Δ*thyA*::*pdeA* were able to grow in the gallbladder (**Fig. 4L**). This suggests that the gallbladder is deficient in pyrimidines, which are necessary to support the growth of Δ*thyA* mutants. This deficiency may contribute to their more pronounced defect in dissemination to the liver and spleen during oral infection compared to retro-orbital infection.

Furthermore, WT::*pdeA* exhibited significantly reduced survival in the gallbladder compared to the WT (**Fig. 4L**), indicating that c-di-AMP may also play a critical role in bile acid resistance.

Taken together, our data indicate that the gallbladder is a significant bottleneck for Δ*thyA* in establishing infection, whereas c-di-AMP is required for Δ*thyA* to colonize the intestines, disseminate from the gastrointestinal tract to peripheral organs, and grow in these organs.

### PstA promotes TLD under low c-di-AMP concentrations

Previous studies have identified the PII family signaling protein PstA as a conserved c-di-AMP binder in Firmicutes (40–42). PII signaling proteins typically regulate cellular processes through protein-protein interactions that are inhibited upon metabolite binding (43). Although the specific targets of PstA remain unknown, deletion of PstA partially suppresses *Lm* cell death under c-di-AMP deprivation in rich medium (33, 41). We hypothesize that PstA plays a role in regulating the bacterial stress response, including thymineless death, downstream of c-di-AMP signaling.

We found that Δ*thyA*::*pdeA* strain exhibited a growth defect compared to Δ*thyA* when grown in low thymidine concentrations. While deletion of *pstA* in the Δ*thyA* background had no effect on growth, its deletion in the Δ*thyA*::*pdeA* strain rescued the growth defect. Furthermore, complementation of *pstA* in the Δ*thyA*Δ*pstA*::*pdeA* strain restored the growth defect to a level similar to that of Δ*thyA*::*pdeA*, further supporting that the growth impairment is attributable to PstA (**Fig. 5A**). Similarly, deletion of *pstA* restored the survival of the Δ*thyA*::*pdeA* strain in the absence of exogenous thymidine (**Fig. 5B**). This phenotype was also observed in the Δ*thyA*Δ*dacA*::*dacA* strain, in which the Δ*dacA* mutation was complemented with an IPTG-inducible *dacA*. In the absence of both thymidine and IPTG, Δ*thyA*Δ*dacA*::*dacA* exhibited a significant survival defect compared to the Δ*thyA* strain, while *pstA* deletion rescued this defect (**Supplementary** Fig. 2). These data indicate that, either overexpression of PdeA or deletion of DacA, both of which reduce c-di-AMP levels, led to a significant cell death of Δ*thyA* in the presence of PstA.

**Figure 5.**
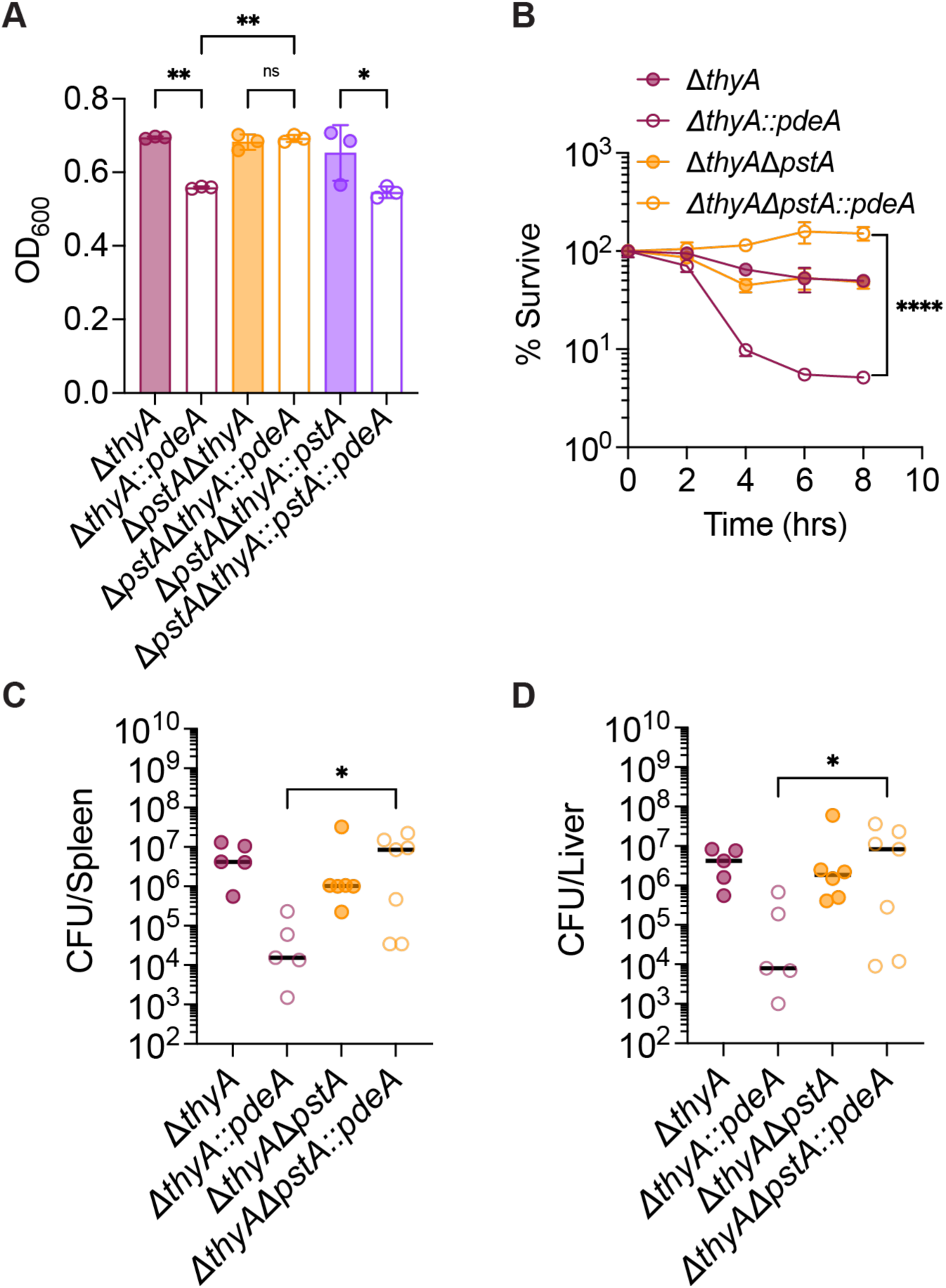
PstA promotes thymineless death in the absence of c-di-AMP. (A) Optical density (600 nm) of *Lm* strains grown in BHI with 1.25 µg/ml thymidine for 6 hours. (B) Bacterial survival in the absence of thymidine. Survival rate at each time point was calculated by normalizing CFU counts to the value at time 0. (C) and (D) CFU recovery of *Lm* from spleens and livers. C57BL/6 mice were infected with 10⁵ CFU *Lm* per mouse via the retro-orbital route. Livers and spleens were harvested for CFU enumeration at 72 hpi. For panels (A) and (B), mean values of biological replicates are plotted, and error bars indicate ±SD*, P* values were calculated using Ordinary one-way ANOVA analysis. For all mouse infection panels, biological replicates are plotted. Horizontal black bars indicate the median of the data. *P* values were calculated using Mann-Whitney analysis. Asterisks indicate that differences are statistically significant (*, *P* < 0.05; **, *P* < 0.01; ***, *P* < 0.001, ****, *P* < 0.0001), and ns indicates no significant difference.

To investigate whether PstA affects the *in vivo* growth of Δ*thyA*, C57BL/6 mice were retro-orbitally infected with Δ*thyA*, Δ*thyA*::*pdeA*, Δ*thyA*Δ*pstA*, and Δ*thyA*Δ*pstA*::*pdeA* strains at an inoculum of 10⁵ CFU per mouse. Consistent with the *in vitro* survival assays, deletion of *pstA* fully restored the growth defect of Δ*thyA*::*pdeA in vivo*, with CFU recovery in both the spleen and liver comparable to that of the Δ*thyA* strain (**Fig. 5C and D**).

Taken together, our data suggest that PstA is detrimental to the survival of Δ*thyA* in the absence of c- di-AMP. Elevated c-di-AMP levels in Δ*thyA* appear to counteract the toxic effect of PstA.

## Discussion

The results of this study demonstrate that c-di-AMP is essential for the survival of *Lm* in response to multiple categories of antibiotics, including β-lactams, antifolates, and protein synthesis inhibitors. Our previous work showed that deactivation of ThyA, either through antifolate treatment or deletion of the *thyA* gene (Δ*thyA*), leads to elevated levels of c-di-AMP (24). In this study, we further demonstrate that the elevated c-di-AMP is necessary to inhibit the toxic activity of PstA and subsequently counteract cell death of Δ*thyA* under thymidine-deprived conditions and during intracellular growth.

TLD underlies the effectiveness of several antibacterial (trimethoprim, sulfamethoxazole), antimalarial (pyrimethamine, sulfonamide), anticancer (methotrexate, fluorouracil), and immune-modulating (methotrexate) agents (16). However, the underlying mechanisms have remained elusive for over 70 years. *Escherichia coli*, the most widely used model for studying bacterial thymineless death, exhibits a brief resistance phase (approximately 1 hour) in the absence of thymidine, during which CFU counts remain stable, followed by a rapid exponential death (RED) phase. This transient resistance is partly attributed to the use of intracellular dTDP-sugars as alternative thymidine sources, which temporarily support chromosomal DNA replication during thymidine starvation (44). In contrast, *Lm* exhibits a significantly prolonged resistance phase lasting up to 8 hours (**Fig. 2D**). *E. coli* does not produce c-di-AMP, whereas *Lm* lacks the ability to utilize dTDP-sugars as precursors for DNA replication. Notably, reducing intracellular c-di-AMP levels in *Lm* Δ*thyA* abolishes the extended resistance phase, leading to a rapid loss of cell viability. These findings suggest that c-di-AMP may regulate an as-yet-unidentified pyrimidine salvage pathway in *Lm* that supplies thymidine during thymidine starvation, thereby sustaining the prolonged resistance phase. Although the mechanism by which c-di-AMP regulates TLD remains to be further studied, we have identified that the conserved c-di-AMP-binding protein PstA, conserved in Firmicutes, serves as a negative regulator of this pathway. Moreover, PstA is found in several human pathogens, including *Lm*, *Staphylococcus aureus*, and *Enterococcus faecalis*. *S. aureus* thymidine auxotrophs, also known as thymidine-dependent small-colony variants, are frequently isolated from cystic fibrosis (CF) patients undergoing antifolate antibiotic treatment and are associated with more severe lung disease worldwide (45–47). Therefore, the role of PstA in regulating TLD may represent a clinically significant, conserved mechanism shared by pathogenic Firmicutes.

PstA belongs to the PII signaling protein family, which are multitasking regulators ubiquitously found across all domains of life. These proteins integrate signals of energy state and carbon/nitrogen balance by binding to various ligands, including nucleotides (ATP, ADP, and AMP), second messengers (cAMP, SAM-AMP), and metabolites (α-ketoglutarate, HCO_3_^-^). Different binding states, along with post-translational modifications (PTMs), influence interactions with a wide range of effector proteins, such as transporters, metabolic enzymes, and transcription factors, thereby modulating various metabolic states of the cell (43). PII proteins contain two flexible loops: the T-loop and the B-loop. PstA is evolutionarily distinct from other PII family members. In PII proteins, the large T-loop mediates protein-protein interactions, whereas in PstA, this loop is shorter and contributes to c-di-AMP binding. Additionally, the B-loop in PstA is larger than in canonical PII proteins and lacks the highly conserved ATP-binding Walker A motif (40, 41). The binding of c-di-AMP induces significant changes in the position and orientation of the B-loop (40, 41), which may impact the interaction between PstA and its binding partners.

Although the downstream binding partners of PstA remain unknown, our findings, along with previous studies, suggest that PstA may act as a multitasking regulator of cell viability under various stress conditions, including antibiotic stress and nutrient starvation, and is detrimental to bacterial survival when c-di-AMP levels are low. *Lm* Δ*dacA* strain, in which c-di-AMP synthesis is abolished, exhibits significantly increased susceptibility to cefuroxime (33). Several studies have shown that deletion of *pstA* in Δ*dacA* restores cefuroxime resistance (33, 48). The bactericidal activity of β-lactam antibiotics under aerobic conditions is driven by the enhanced glycolytic flux they induce, which leads to an increased generation of ROS from the respiratory chain, ultimately resulting in cell death (49). Furthermore, the PstA-mediated increase in β- lactam susceptibility depends on the respiratory electron transport chain and is prominent only under aerobic conditions, with the effect largely diminished under hypoxia (48). These data suggest that the presence of PstA may lead to metabolic imbalance when c-di-AMP levels are low, which in turn impacts β-lactam susceptibility. Furthermore, Δ*dacA* exhibits increased cell lysis when grown in rich medium due to (p)ppGpp accumulation, which leads to metabolic imbalance and subsequent inactivation of the pleiotropic transcriptional regulator CodY (50), and depletion of *pstA* partially rescued this cell death (33, 41).

Our study demonstrated that the Δ*thyA* exhibits significant intracellular growth defects, which lead to reduced bacterial burden in both murine intravenous and oral infection models. However, the growth defect was more pronounced following oral infection. In intravenous infections, *Lm* rapidly colonizes the liver, which serves as a major reservoir for dissemination(51). In contrast, during oral infection, *Lm* disseminates from the gastrointestinal tract to peripheral organs, including the gallbladder, which becomes one of the major primary bacterial reservoirs (39). Notably, our study found that Δ*thyA* exhibited a comparable ability to colonize the gastrointestinal tract as the WT strain. However, Δ*thyA* was unable to survive in the gallbladder. Similarly, defects in purine biosynthesis led to severely attenuated growth of *Lm* in *ex vivo* gallbladders and during intragastric infections (39, 52, 53). These findings suggest that bile may be limited in either pyrimidine or purine availability, or that access to these nucleotides is restricted in this environment. In summary, our study revealed a previously unrecognized role of c-di-AMP in modulating bacterial thymineless death. We anticipate that these findings will provide valuable insights into the physiological functions of the widespread bacterial second messenger c-di-AMP and broaden our understanding of its impact on antifolate antibiotic resistance and treatment outcomes in c-di-AMP-producing bacteria.

## Materials and Methods

### Microbe strains and culturing conditions

*Escherichia coli* strains used for cloning were grown in Lysogeny Broth (LB) at 37°C with shaking. *Lm* mutants were derived from the WT strain 10403S (54). All *Lm* strains were cultured statically in Brain Heart Infusion (BHI) broth at 30°C unless otherwise stated. Antibiotics were used at the following concentrations: streptomycin, 200 µg/mL; chloramphenicol, 50 µg/mL (*E. coli*), 10 µg/mL (*Lm*); tetracycline, 2 µg/mL; ampicillin, 100 µg/mL; kanamycin, 50 µg/mL; nalidixic acid 25 μg/mL.

### Plasmid and strain construction

The strains used in this study are listed in **Table S1**, and the primers are listed in **Table S2**. All *Lm* knockout mutants were generated by allelic exchange using the conjugation-proficient plasmid PliM, which expresses a mutated phenylalanyl-tRNA synthetase gene (*pheS\**). PheS*\** incorporates the toxic DL-4- chlorophenylalanine into cellular proteins, leading to bacterial cell death and enabling counterselection(55). Briefly, the PliM vector carrying 500 bp of upstream and downstream arms of the target gene was transferred to *E. coli* SM10 cells. The SM10 strain was mixed with the *Lm* strain and incubated for 6 hours on BHI agar plates without antibiotics at 30°C for conjugation. Afterward, selection was performed on BHI plates containing both streptomycin and chloramphenicol to select against the *E. coli* and plasmid-free *Lm*. Single *Lm* colonies were passaged three times on BHI plates with streptomycin and chloramphenicol at 42°C to ensure plasmid integration. Single colonies were then passaged twice on BHI plates with 18 mM DL-4-chlorophenylalanine for allelic exchange. Overexpression strains were generated by integrating the pPL2 plasmid (56) into the *Lm* chromosome under the control of the p*Hyper* promoter (57) through conjugation, followed by selection on BHI plates supplemented with tetracycline. Complementation strains were generated by electroporating the pBAV1K-E plasmid (58) backbone carrying the genes under their native promoters, followed by selection on BHI plates supplemented with kanamycin. The resulting mutants were verified through Sanger sequencing and further confirmed by qRT-PCR.

### Antibiotic susceptibility assays

For the antibiotic susceptibility screenings, the mid-exponential phase of *Lm* cultures was diluted with fresh BHI to an OD_600_ of 0.05. A 50 μL aliquot of the bacterial suspension was added to the BIOLOG^TM^ microplates (PM11C and PM12B), sealed with an oxygen-permeable film, and incubated statically at 30°C for 6 hours. The bacterial suspension was then serially diluted and plated on BHI plates for CFU enumeration.

For the disk diffusion assays, 10 μL of overnight *Lm* culture was added to 3 mL of melted top agar (0.8% agar in 0.8% NaCl solution, pre-cooled to 55°C), mixed by inversion, and then poured onto warm BHI agar plates. Once the plates had solidified, a 7 mm × 3 mm Whatman paper disk was placed on the top agar, and 5 μL of antibiotic solutions were applied to each disk. The plates were incubated at 30°C until the inhibition zone appeared.

### Bacterial growth and survival assays

The mid-exponential phase of *Lm* cultures was washed twice with PBS and pelleted by centrifugation. For the bacterial growth curve, the *Lm* pellet was resuspended in BHI to an OD₆₀₀ of 0.05 with various concentrations of thymidine. A 200 μL aliquot of the bacterial suspension was added to 96-well suspension plates sealed with oxygen-permeable film and cultured statically at 30°C. The OD₆₀₀ was measured every 30 minutes using a Microplate Reader (BioTek). For bacterial survival assays under thymidine starvation, the *Lm* pellet was resuspended in LB to an OD₆₀₀ of 0.1 and cultured at 37°C with shaking. At various time points, the bacterial culture was serially diluted and plated on BHI plates supplemented with thymidine for CFU enumeration.

### Quantification of c-di-AMP

The intracellular c-di-AMP of *Lm* strains was extracted using methanol and quantified using a luminescence-based coupled enzyme assay for c-di-AMP quantification (CDA-Luc), as previously described (59). Briefly, *Lm* at the mid-exponential phase, grown in BHI under the indicated conditions, was measured for OD₆₀₀ and plated for CFU enumeration. A total of 2 mL of bacterial culture was collected by centrifugation (15,000 × g, 5 min), resuspended in 50 μL of Milli-Q water to form a homogeneous suspension, and 500 μL of methanol was added. The bacteria were lysed by sonication at a 40% amplitude setting for 5 pulses (1 second on/1 second off) using a sonicator (Fisher Scientific, USA). The methanol extract was collected by centrifugation, and the bacterial pellet was washed with an additional 500 μL of methanol. Both methanol extracts were combined and dried at room temperature under the “V-AL” setting of a Vacufuge Concentrator (Eppendorf, USA) with caps open to allow complete evaporation of methanol. Each sample was then resuspended in assay buffer for c-di-AMP quantification using the CDA-Luc assay (59).

### Macrophage infections

For the study of intracellular growth of *Lm*, 0.5 × 10⁶ iBMDMs were plated in 24-well tissue culture plates and incubated overnight at 37 °C with 5% CO₂. Mid-exponential phase *Lm* cultures grown in BHI containing 10 µg/mL thymidine were diluted 2,500- to 3,000-fold in BMM medium (DMEM GlutaMAX supplemented with 10% FBS and 10% L929 supernatants) to standardize bacterial concentrations. Macrophages were incubated with the *Lm* suspension at a multiplicity of infection (MOI) of 1. After a 0.5 hr incubation, the macrophages were washed once with PBS and given fresh BMM medium containing 50 µg/mL gentamicin to eliminate extracellular bacteria. At various time points post-infection, macrophages were washed once with PBS and lysed by adding H₂O, followed by incubation at 37 °C for 5 minutes. Appropriate dilutions of the lysate were plated on BHI agar and incubated overnight at 37 °C for *Lm* CFU enumeration. For the exogenous thymidine supplementation experiment, 100 µg/mL thymidine was added to the *Lm* suspension at the time of infection and maintained in the BMM medium throughout the experiment.

### Plaque assays

Murine L2 fibroblast cells were cultured in BMM medium (DMEM GlutaMAX supplemented with 10% fetal bovine serum). Plaque assays were performed on L2 fibroblast monolayers as previously described(37), with the following modifications. A total of 1.2 × 10⁶ cells were seeded in 6-well tissue culture plates in 3 mL of BMM medium and grown overnight to form monolayers. The L2 monolayers were infected with *Lm* strains at an MOI of 0.2. After 1 hour of incubation, extracellular *Lm* cells were washed off with PBS, and BMM medium containing 0.7% Superpure agarose (#G02PD-125), 10 µg/mL gentamicin, and varying concentrations of thymidine was preheated to 56°C and added to the cells. At 2 days post-infection, the staining solution (BMM medium containing 0.7% agarose and 0.25% neutral red) was added and incubated for 18 hours before the plates were scanned and plaques were analyzed using ImageJ.

### Microscopic analysis

For the study of intracellular *Lm* morphology, 10^5^ iBMDMs were plated in 96-well poly-D-lysine-coated glass-bottom plates (MatTek Life Sciences, P96GC-1.5-5-F) and incubated overnight at 37°C with 5% CO₂. Macrophages were infected with *Lm* at a multiplicity of infection (MOI) of 10 or 20. After a 0.5-hour incubation, the macrophages were washed once with PBS and given fresh BMM medium containing 50 µg/mL gentamicin to eliminate extracellular bacteria. At various time points post-infection, macrophages were washed twice with PBS and fixed in 4% paraformaldehyde in PBS for 15 minutes at room temperature.

The cells were washed in Tris-buffered saline (TBS) with 0.1% Triton X-100 and blocked in TBS with 1% bovine serum albumin (BSA) at room temperature for 1 h. The *Lm* was stained with rabbit anti-*Lm* antibody (Abnova, PAB29797) at a 1:200 dilution in TBS with 1% BSA overnight at 4°C. The macrophages were washed five times with 200 µL TBST buffer and incubated with Alexa Fluor 488-conjugated goat anti-rabbit IgG secondary antibody (Invitrogen, A11008) at a 1:200 dilution, Alexa Fluor 568 phalloidin (Invitrogen, A12380) at a 1:1000 dilution, and DAPI at a 1:1000 dilution in TBS with 1% BSA for 2 hours at 4°C. The cells were then washed five times with 200 µL TBST buffer, maintained in TBS, and visualized under oil immersion at 100× using a Nikon Eclipse fluorescence microscope.

For the live/dead staining of bacteria, mid-exponential phase *Lm* cultures were washed twice with sterile PBS and diluted with fresh LB to an OD_600_ of 0.1, supplemented with 0.1 mg/mL propidium iodide. A 5 μL aliquot of the bacterial suspension was placed onto an agarose gel pad (1% agarose in LB broth) an Propidium Iodide d air-dried. The agar pad was then covered with a coverslip and sealed with glue. The bacteria were visualized under oil immersion at 100× magnification using a Nikon Eclipse microscope.

### Mouse infection

All the mice used in this study were of the C57BL/6J background and purchased from Jackson Laboratories. The mice were maintained under SPF conditions, as ensured by the rodent health monitoring program overseen by the Animal Care Facility of the Shimadzu Institute for Research Technologies at UT Arlington. All experiments involving mice were performed in compliance with the guidelines set by the American Association for Laboratory Animal Science (AALAS) and were approved by the Institutional Animal Care and Use Committee (IACUC) at UT Arlington. All experiments were carried out using mice aged 7-10 weeks, matched by gender, age, and body weight.

Prior to animal experiments, *Lm* strains were cultured overnight statically at 30°C in BHI, then back diluted 1:5 in fresh BHI and grown for 1 hour at 37°C with shaking. The bacterial suspension was diluted with cold PBS. For systematic infection, a 200 μL aliquot of the *Lm* suspension was retro-orbitally injected into the mice with an inoculum of 10^5^ CFU/mouse. For the oral infection, mice will be given 5 mg/mL streptomycin in water with regular food for 48 hours, followed by a 16-hour fasting. The mice were orally instilled with 20 μL of *Lm* suspension in PBS via pipette, with an inoculum of 10^8^ CFU/mouse. After infection, the infected mice will be provided with regular food and water. At 72 hpi, the mice from both infection models were euthanized by CO_2_, and the liver, spleen, gallbladder, cecum, small intestine, colon, and feces were collected. The liver, spleen, cecum, small intestine, and colon were homogenized and lysed in 10 mL of lysis buffer (0.1% IPEGAL in water) using a Tissue Tearor Homogenizer. The gallbladder was opened using a sterile wooden stick in 500 μL of lysis buffer, and fecal samples were mixed in 1 mL of lysis buffer by vortexing. Bacterial burdens were enumerated by plating serial dilutions on BHI plates containing streptomycin and 10 μg/mL thymidine to support the growth of Δ*thyA* mutants. For intestinal and fecal samples, an additional 25 μg/mL nalidixic acid was added to prevent contamination from the microbiota.

### Quantification and statistical analysis

All numerical data were statistically analyzed and plotted using GraphPad Prism software. For bacterial growth curves, survival assays, tissue culture experiments, and c-di-AMP quantification plots, mean values of biological replicates are plotted, with error bars indicating ±SD. Data shown in animal infection plots are represented as the median of biological replicates. The exact numbers of replicates are stated in the figure legends. Statistical tests used and the corresponding statistical parameters for each experiment are detailed in their respective figure legends. No methods were employed to determine whether the data met the assumptions of the statistical approach.

## Acknowledgments

We would like to thank Joshua J. Woodward of University of Washington for providing iBMDMs, murine L2 fibroblast cells, Δ*dacA*::*dacA Lm* strain, and experimental and intellectual support. We thank members of the Boutte lab at the UT Arlington for valuable discussion and feedback. We also thank the Boutte, Pellegrino, and Ghose labs at UT Arlington for reagents and equipment sharing. This work was supported by the National Institute of Allergy and Infectious Diseases (R01AI116669, J.J.W.) and the UT System STARs (Science and Technology Acquisition and Retention) program (Q.T.).

## Author contributions

Q.T. and J.J.W. conceived and designed the research. Q.T., J.P.L., O.M.E., and D.O. performed experiments.

Q.T. and J.J.W. provided key insights. J.J.W. provided key tools and reagents. Q.T., J.P.L., and O.M.E. analyzed the data and wrote the paper. All authors reviewed the manuscript prior to submission.

## Appendixes

**Supplemental Table 1.** Strains used in this study.

| Strain Name | Genotype | Background | Plasmid | Reference |
| --- | --- | --- | --- | --- |
| TQ572 | WT | *L. monocytogenes* 10403S |  | (54) |
| TQ290 | WT::*pdeA* | *L. monocytogenes* 10403S | pPL2-*pdeA* | This work |
| TQ268 | Δ*thyA* | *L. monocytogenes* 10403S | | This work |
| TQ288 | Δ*thyA*::*pdeA* | *L. monocytogenes* 10403S | pPL2-*pdeA* | This work |
| TQ322 | Δ*thyA*::*pdeA*<sub>84-657</sub> | *L. monocytogenes* 10403S | pPL2-*pdeA*<sub>84-657</sub> | This work |
| TQ326 | Δ*thyA*::*pdeA*<sub>DHH-AAA</sub> | *L. monocytogenes* 10403S | pPL2-*pdeA*<sub>DHH-AAA</sub> | This work |
| TQ304 | Δ*thyA*Δ*pstA* | *L. monocytogenes* 10403S | | This work |
| TQ306 | Δ*thyA*Δ*pstA*::*pdeA* | *L. monocytogenes* 10403S | pPL2-*pdeA* | This work |
| TQ452 | Δ*dacA*::*disA* | *L. monocytogenes* 10403S | pLIV2-*disA* | (32) |
| TQ446 | Δ*dacA*::*dacA* | *L. monocytogenes* 10403S | pLIV2-*dacA* | (32) |
| TQ449 | Δ*dacA*::*dacA* Δ*pstA* | *L. monocytogenes* 10403S | pLIV2-*dacA* | This work |
| TQ282 | Δ*dacA*::*dacA* Δ*thyA* | *L. monocytogenes* 10403S | pLIV2-*dacA* | This work |
| TQ305 | Δ*dacA*::*dacA* Δ*thyA*Δ*pstA* | *L. monocytogenes* 10403S | pLIV2-*dacA* | This work |
| TQ444 | SM10 | *E. coli* SM10 |  |  |
| TQ443 | XL1B | *E. coli* XL1Blue |  |  |

**Supplemental Table 2.**
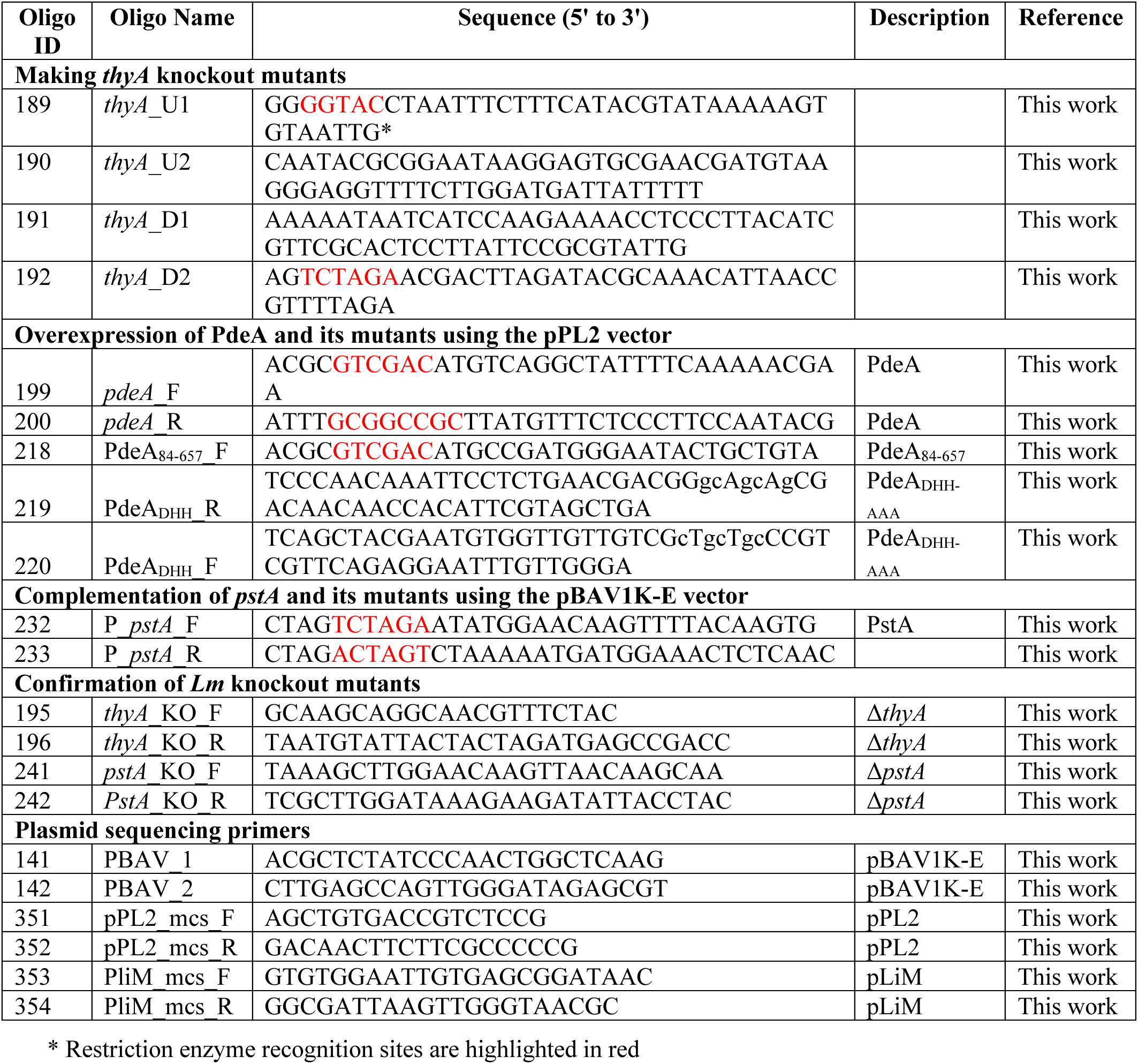
Oligonucleotides used in this study.

**Supplemental Fig. 1.**
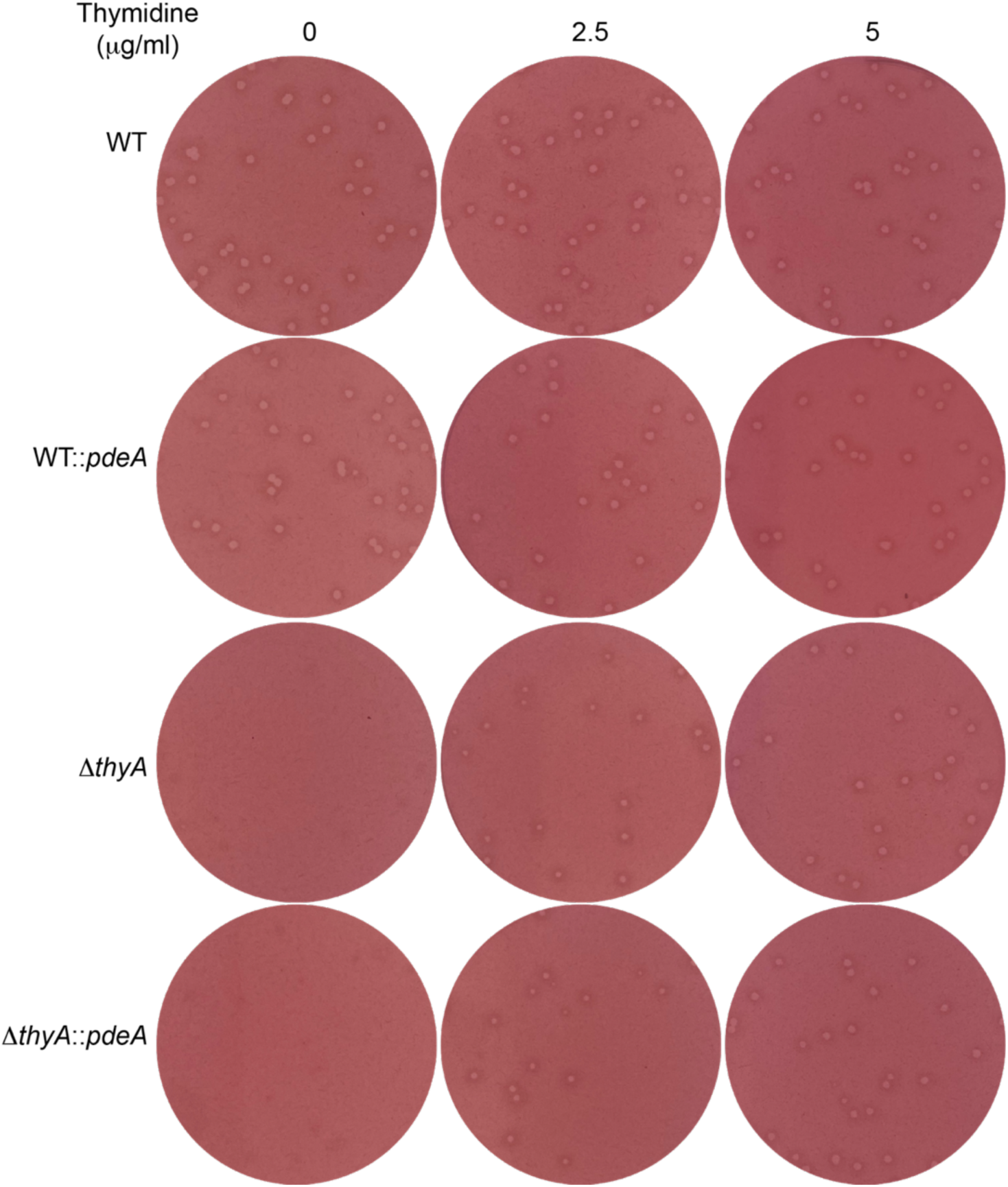
Plaques in L2 fibroblasts infected with WT, WT::*pdeA*, Δ*thyA*, and Δ*thyA*::*pdeA* strain strains. The L2 monolayers were infected with *Lm* strains at an MOI of 0.2. After 1 hour of incubation, extracellular *Lm* cells were washed off with PBS, and BMM medium containing 0.7% Superpure agarose (#G02PD-125), 10 µg/mL gentamicin, and varying concentrations of thymidine was preheated to 56°C and added to the cells. At 2 days post-infection, the staining solution (BMM medium containing 0.7% agarose and 0.25% neutral red) was added and incubated for 18 hours before the plates were scanned.

**Supplemental Fig. 2.**
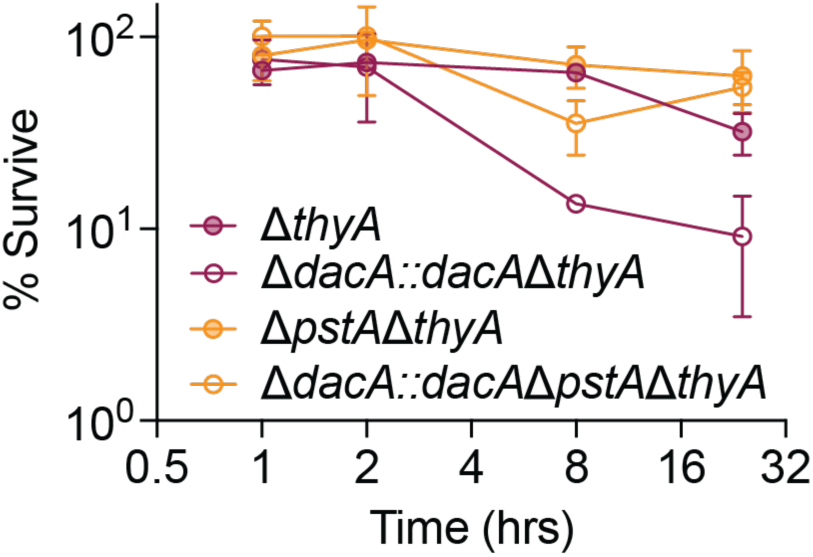
Bacterial survival in the absence of thymidine. The percentage of survival at each time point was calculated by normalizing the CFU count to the value at time 0.

